# Enhanced Diffusion through Multivalency

**DOI:** 10.1101/2023.09.20.558647

**Authors:** Ladislav Bartoš, Mikael Lund, Robert Vácha

## Abstract

In multivalent systems, multiple ligands from one entity simultaneously bind to multiple receptors on another entity. These interactions are of crucial significance in a wide range of biological and technological mechanisms, encompassing selectivity, host recognition, viral penetration, therapeutic delivery, as well as the adhesion phenomena found in cells, polymers, and nanoparticles. In this study, we used computer simulations to investigate 1D and 2D diffusion of adsorbed particles with varying valency but with the same overall affinity to the host. We demonstrate a remarkable diffusion acceleration for particles with increasing valency. Non-diffusing monovalent particle can attain almost unrestricted diffusion when becoming multivalent while retaining its affinity for the host tether or surface. Moreover, diffusion of multivalent particles with rigid ligand distribution can be controlled by patterned host receptors. Our results have practical implications for the design of fast-diffusing particles that maintain a strong affinity for target surfaces or molecules.

**TOC Graphic:** 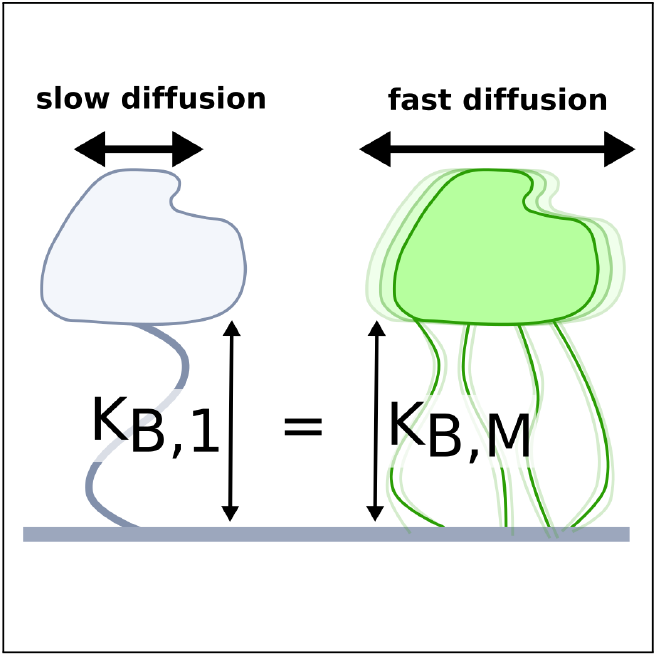

**Significance:** We investigated how the number of binding sites (referred to as valency) on particles or entities impacts their movement when attached to surfaces or filaments. Valency can be understood as how many “hands” a particle has to grip the surface. Surprisingly, particles with more “hands” move faster if they hold onto the surface with the same strength. Furthermore, the motion of these particles can be controlled by designing surfaces with specific patterns that the “hands” can grasp. This means that we can design particles that move rapidly while remaining attached to the desired locations. These findings hold promise for applications like drug delivery and materials technology, and for understanding biological processes.

## Introduction

Multivalent binding refers to the simultaneous non-covalent interaction of one entity with multiple binding ligands on another entity. This mode of interaction is frequent in biological systems, involving diverse entities such as proteins,^1–3^ antibodies,^4^ carbohydrates,^5–7^ and viruses,^8–10^ but also in colloids, polymers, and nanoparticles.^11–13^

The unique characteristic of multivalent binding lies in its ability to amplify the strength and specificity of molecular recognition. By engaging multiple receptor sites, a molecule can enhance its binding affinity and selectivity towards its target,^14,15^ enabling biological processes such as cell adhesion, ^16,17^ signaling, ^18,19^ immune response modulation,^20,21^ or viral entry.^8^ Furthermore, multivalent molecules encompass molecular walkers *such as* kinesin,^22^ myosin V,^23^ and various synthetic counterparts.^24^ Molecular walkers possess the ability to traverse fibers or surfaces while maintaining their interaction without complete detachment. This characteristic renders them not only significant in a biological contexts, but also holds promise for applications in other fields such as medicine or nanotechnology.^25,26^

In this study, we present a simple numeri-cal model of particles diffusing/walking on host surfaces or tethers to provide the relationship between the valency of the particles and their diffusion rates. We restrict the study to particles with identical affinity towards the host, and demonstrate that increased valency dramatically accelerates diffusion. In addition, we show the impact of the rigid distribution/geometry of host receptors on the diffusion rates of the particles with matching distribution of ligands. Our findings provide a foundation for the design of particles with controlled diffusion capabilities on the 1D or 2D target.

## Methods

The numerical model consisted of *N* diffusing ligands connected by linkers represented by harmonic springs with force constant *k* and equilibrium distance *λ*_eq_, the later defining the unit length. In the following text, we refer to a group of linked ligands as a particle. All particles were able to translate either on a line (1D) or on a plane (2D) decorated with an oscillating potential given by

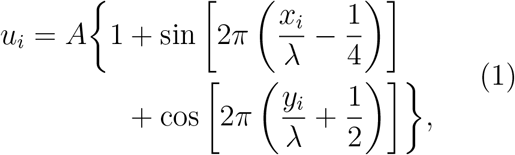

where *{x*_*i*_, *y*_*i*_*}* is the *i*’th ligand position (for 1D, *y*_*i*_ = 0), *A* is the amplitude, *λ* is the distance between the potential wells (see Figure 1). The potential wells represented the receptor sites on the host surface and the amplitude (*A*) determined the affinity of the ligand to the receptors. Only *n ≤N* of the *N* ligands interacted with the potential whereby the net surface affinity was set *K*_*B*_ = *An* for varying *n*. Two connectivity patterns between the ligands were simulated: a linear chain and a star, where all ligands were connected to one central ligand (see Figure 1). See Tables S1 and S2 for detailed information about the simulated patterns.

**Figure 1:**
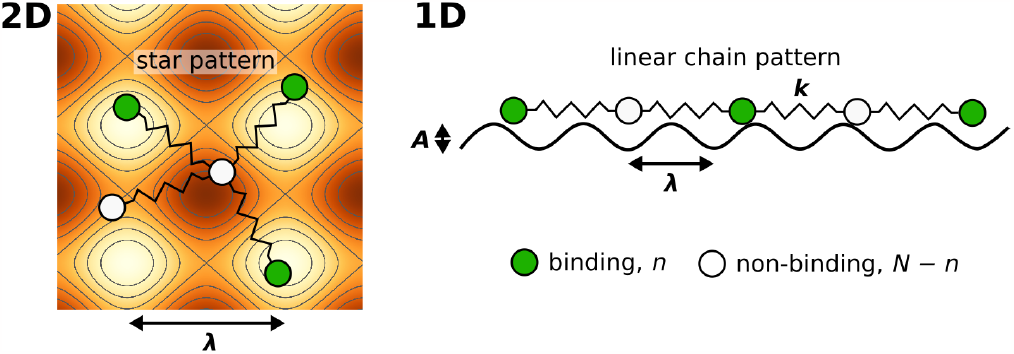
Illustration of the employed model. The diffusing particle was composed of *N* ligands connected via linkers of specific stiffness *k. n* of the ligands were binding, i.e. interacting with the surface potential, which corresponds to the valency of the particle. In the 2D case, the surface potential represented a host surface such as a cell membrane with uniformly distributed receptors. In the 1D case, the potential represented a tether such as a microtubule or a DNA double helix. The distance between the receptors was specified by the parameter *λ* while the affinity of the particle to the surface was set by the parameter *K*_*B*_ = *An*.

The system was propagated using Dynamic Monte Carlo (MC) where individual ligands were moved using MC translation moves, accepted or rejected according to MetropolisHastings criterion. ^27^ The maximum displacement was arbitrarily set to 0.1*λ*_*eq*_ which also defined the time-scale of the simulation. The code used to perform the simulations is available from doi.org/10.5281/zenodo.8340209.

Initially, every replica of each system was equilibrated for either 50,000 (most systems) or 200,000 (systems with *N* = 10, *n* = 1, and *k ≤*10) sweeps, followed by 30,000 production sweeps. For each sweep, *N* translational attempts were performed. Every 100 sweeps, we calculated the mean-square deviation (MSD) between the current center of geometry of the diffusing ligands and their center of geometry at the start of the simulation. 3000 of these independent replicas were used to collect MSD data and construct a single MSD curve. A line was then fitted through the production part of the collected MSD curve and from the slope of the fitted line, we obtained an estimate of the diffusion rate (diffusion coefficient). We repeated the entire process of collecting MSD data 10 times and averaged the calculated diffusion rates. The error was taken as a standard deviation of the calculated diffusion rates. See Figure S1 for an example of MSD curves calculated for a selected system. In total, each system was simulated with 30,000 independent replicas, resulting in 2.4 (most systems) or 6.9 (*N* = 10, *n* = 1, *k ≤*10) billion MC sweeps being performed for each system. The calculated diffusion rates, *D*, were normalized with the corresponding free diffusion, *D*_0_, in the absence of any surface potential. The obtained relative diffusion rate *D/D*_0_ thus depicts how much slower the diffusion of the bound particle was compared to the diffusion of free particle.

## Results

We began by examining the impact of multiple *weakly* binding ligands compared to a single *strongly* binding ligand. Figure 2 displays the diffusion of particles composed of five ligands that form either a linear chain (1D) or a star (2D). Each particle had a different valency, *n*, i.e. a varying number of binding ligands. We calculated diffusion rates for particles with three different net host surface affinities (binding constants), *K*_*B*_, which remained the same regardless of *n*. The diffusion rates are displayed as relative values to the unbound particles, i.e. free diffusion in the absence of surface potential.

**Figure 2:**
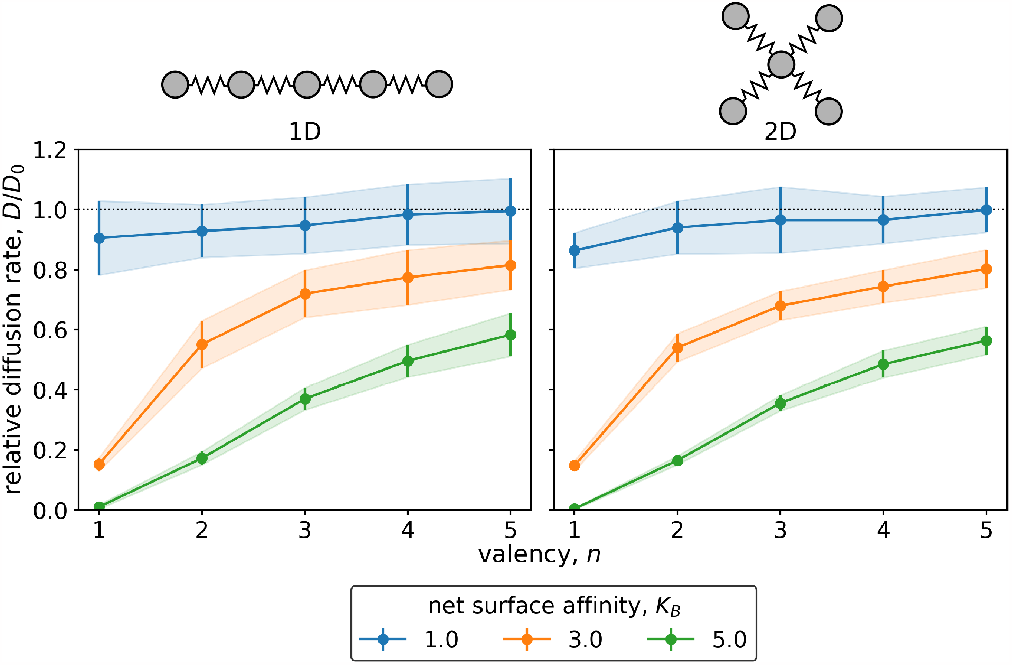
The diffusion of particles comprised of five ligands (*N* = 5) with varying valency, i.e. the number of binding ligands, *n*, ranging from 1 to 5, and three different net surface affinities, *K*_*B*_. Linkers represented by harmonic bonds with a force constant of *k* = 1 connected the ligands, with the distance between the potential wells of the surface equivalent to the equilibrium linker lengths. The relative diffusion rates, *D/D*_0_, represent the diffusion of bound particle with respect to freely diffusing particle without any surface potential. The left plot depicts the diffusion of a linear chain on a line, while the right plot is for star arrangement of ligands diffusing on a plane.

As expected, particles with stronger net surface affinity, *K*_*B*_, diffused less than those with weaker net affinity. (Specifically, particles with *K*_*B*_ = 5 exhibited lower diffusion rates than particles with *K*_*B*_ = 1.)

The relative diffusion rate was dramatically affected by valency, *n*. Increasing the number of binding ligands led to faster diffusion for all surface affinities when compared to the particle with only one binding ligand (*n* = 1). Note that with increasing valency the diffusion rate seems to be converging to free diffusion, *D*_0_.

The correlation between the valency, *n*, and the relative diffusion rate applied to all studied net surface affinities, as demonstrated in Figure 3. This relationship was particularly pronounced for particles containing ten ligands, where the surface affinity per ligand, *A* = *K*_*B*_*/n*, could decrease significantly more than for a 5-ligand particles. For 10-ligand particles with flexible linkers (*k* = 1), it was possible to increase the diffusion rate from nearly immobile (*D/D*_0_ *<* 0.1) to almost as rapid as a non-binding particle (*D/D*_0_ *≈*0.9) by redistributing the surface affinity from one to ten binding ligands. Note again that this redistribution did not impact the net affinity (binding free energy) towards the surface.

**Figure 3:**
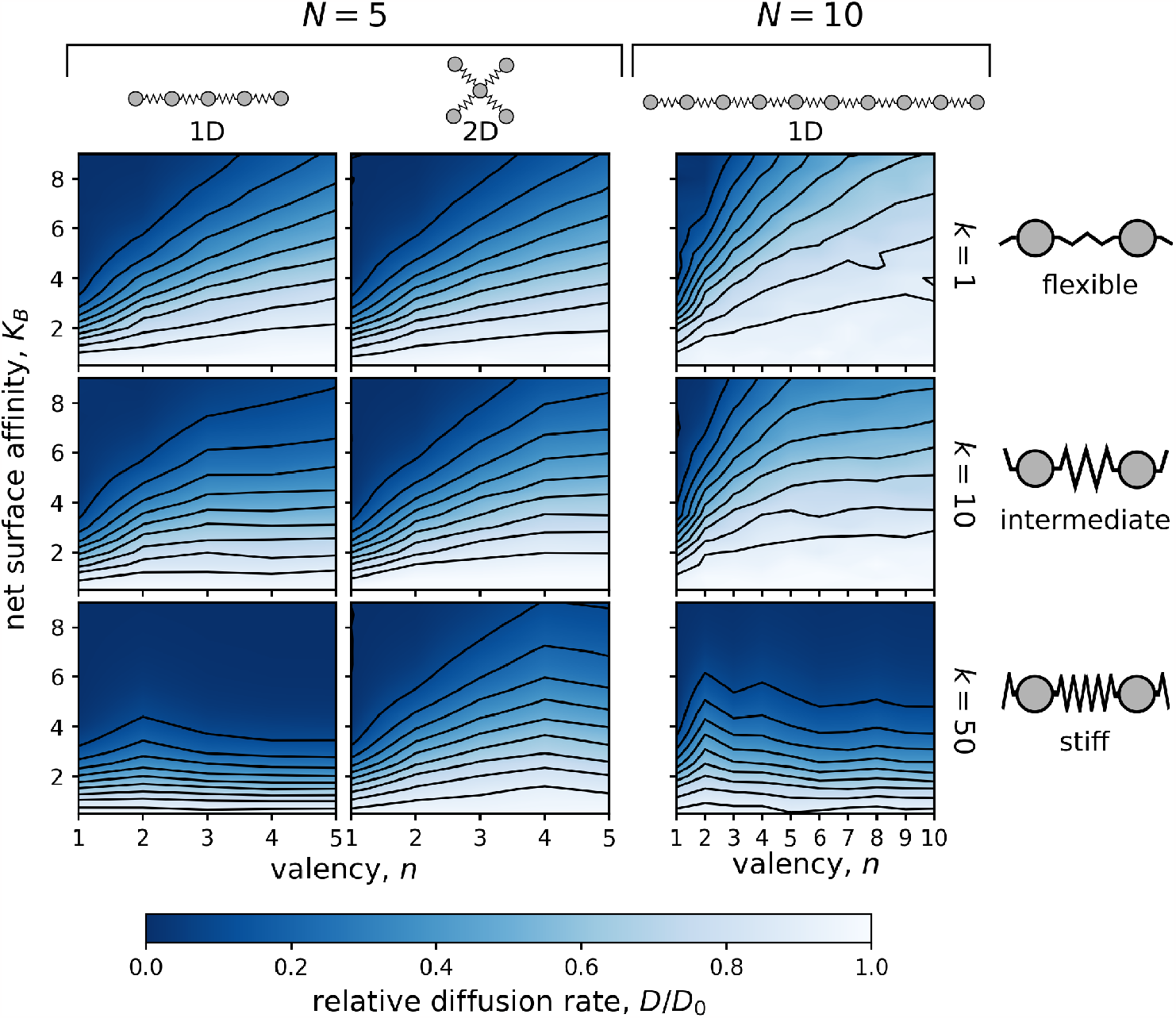
Dependence of the relative diffusion rate, *D/D*_0_, on the valency, *n*, and the net surface affinity, *K*_*B*_. Simulations were performed on particles consisting of either *N* = 5 (two columns on the left) or *N* = 10 ligands (column on the right) connected by flexible (*k* = 1; top row), intermediate (*k* = 10; middle row), or stiff (*k* = 50; bottom row) linkers. The distance between the surface potential wells matched the equilibrium linker lengths, i.e. *λ/λ*_eq_ = 1.0. Diffusion of particles with *N* = 5 was calculated in both 1D (linear chain) and 2D (star-shaped) geometries, while diffusion of particles with *N* = 10 was examined only in the 1D case (linear chain).

In the 1D case, the impact of valency on diffusion was diminished by increasing the stiffness of the linkers connecting the ligands of the particle. In fact, when the force constant of the linkers, *k*, was increased to 50, the diffusion rate became nearly independent of the valency for both *N* = 5 and *N* = 10 particles. The use of stiff linkers resulted in an almost rigid particle with evenly distributed ligands. The spacing between the ligands matched the distance between the surface potential wells. With this “pattern matching”, all binding ligands had to traverse a potential barrier simultaneously leading to the equal height of a diffusion barrier independent of the valency. A similar, but much weaker, effect was also observed for the 2D case.

To examine the “pattern matching” effect in more detail, we investigated the relationship between the equilibrium linker length and the potential wells distance and their effect on the diffusion rate. We assessed the diffusion rates of particles with *N* = 5 and *n* = 5 in both 1D and 2D space for various distances between the potential wells. As depicted in Figure 4, particles with flexible linkers (*k* = 1) were not significantly affected by the altered distance between the potential wells.

**Figure 4:**
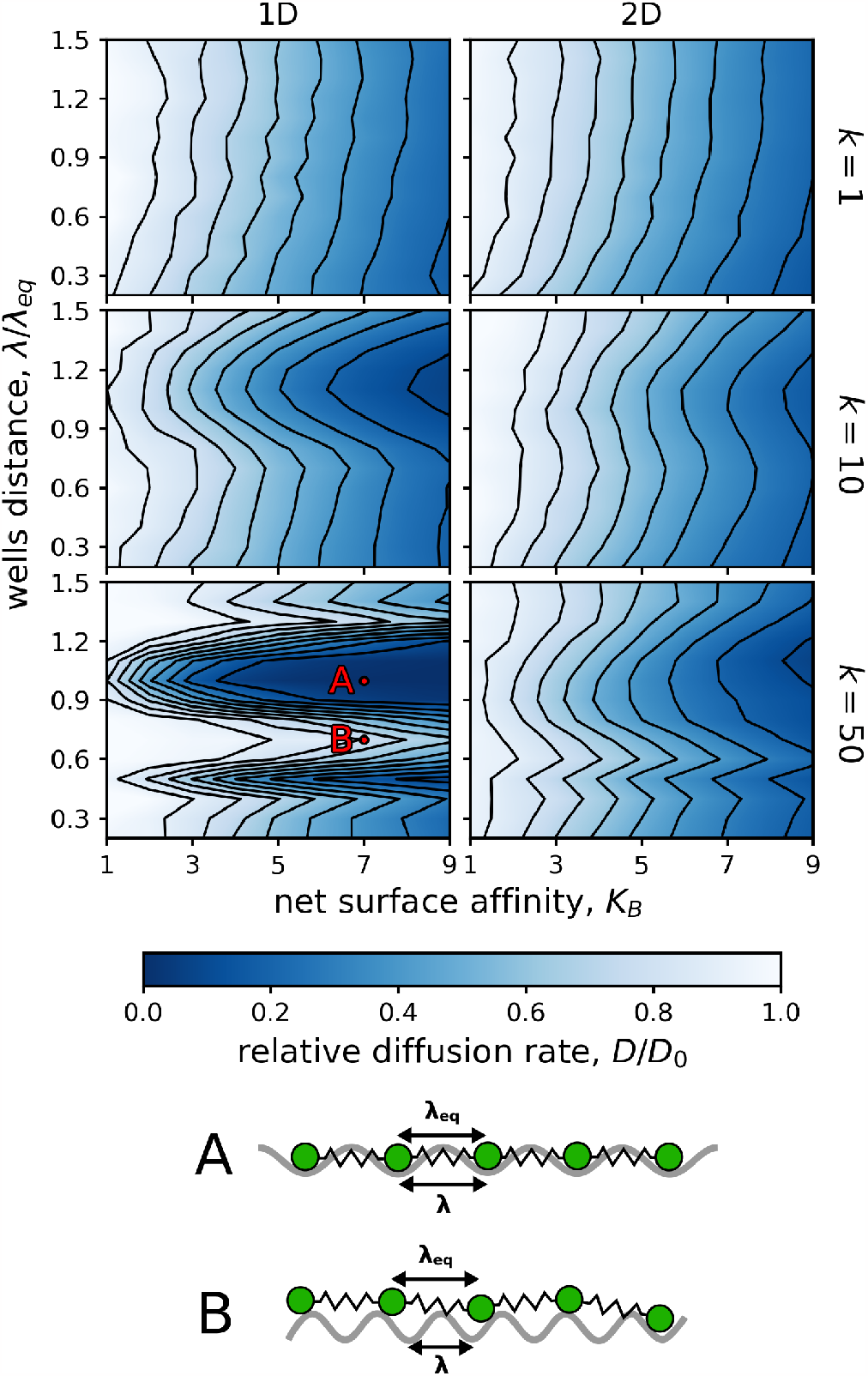
Dependence of the relative diffusion rate, *D/D*_0_, on the net surface affinity, *K*_*B*_, and the distance between the potential wells, *λ/λ*_eq_. The particles were composed of five binding ligands (*N* = 5, *n* = 5), which were connected by flexible (*k* = 1; top row), intermediate (*k* = 10; middle row), or stiff (*k* = 50; bottom row) linkers. The ligands were arranged either as a linear chain (1D) or as a star (2D). A. Schematic representation of a linear chain which ligands are all positioned in the potential surface wells as *λ/λ*_eq_ = 1.0. In such case, diffusion is not enhanced with the increasing valency. B. Schematic representation of a linear chain with linkers not matching the potential surface pattern as *λ/λ*_eq_ = 0.7. In such case, the diffusion is enhanced with increasing valency.

In contrast, the diffusion of particles with intermediately flexible linkers (*k* = 10) was significantly reduced when the distance between the wells of the surface potential roughly matched the equilibrium length of the linkers connecting the ligands.

Interestingly, the relative diffusion rate had its minimum at a distance of *λ/λ*_eq_ = 1.1, instead of the anticipated 1.0, indicating that the particle diffused the slowest when the distance between the potential wells was slightly *longer* than the equilibrium linker length. The origin of this effect is entropic and relates to the system setup, further information is in the Supporting Information.

Particles with stiff linkers (*k* = 50) showed a significant reduction in diffusion not only at wells distances matching the linker lengths but also at *λ/λ*_eq_ *≈*0.5, which was especially noticeable in the 1D case. This behavior was not surprising since at this well spacing, all binding ligands also need to be simultaneously at the potential barriers for further diffusion. Furthermore, in Figure 5, we show that increasing the valency of particles with stiff linkers (*k* = 50) had almost no impact on the relative diffusion rate if the equilibrium linker length roughly matched the wells distance. This is in line with the same effect in Figure 3. Moreover, the effect of “pattern matching” can be modulated by using a surface potential with “nonmatching” wells distances. For instance, *λ/λ*_eq_ = 0.7 can lead to *increased* effect of valency for stiffly linked ligands, as shown in Figure S2.

**Figure 5:**
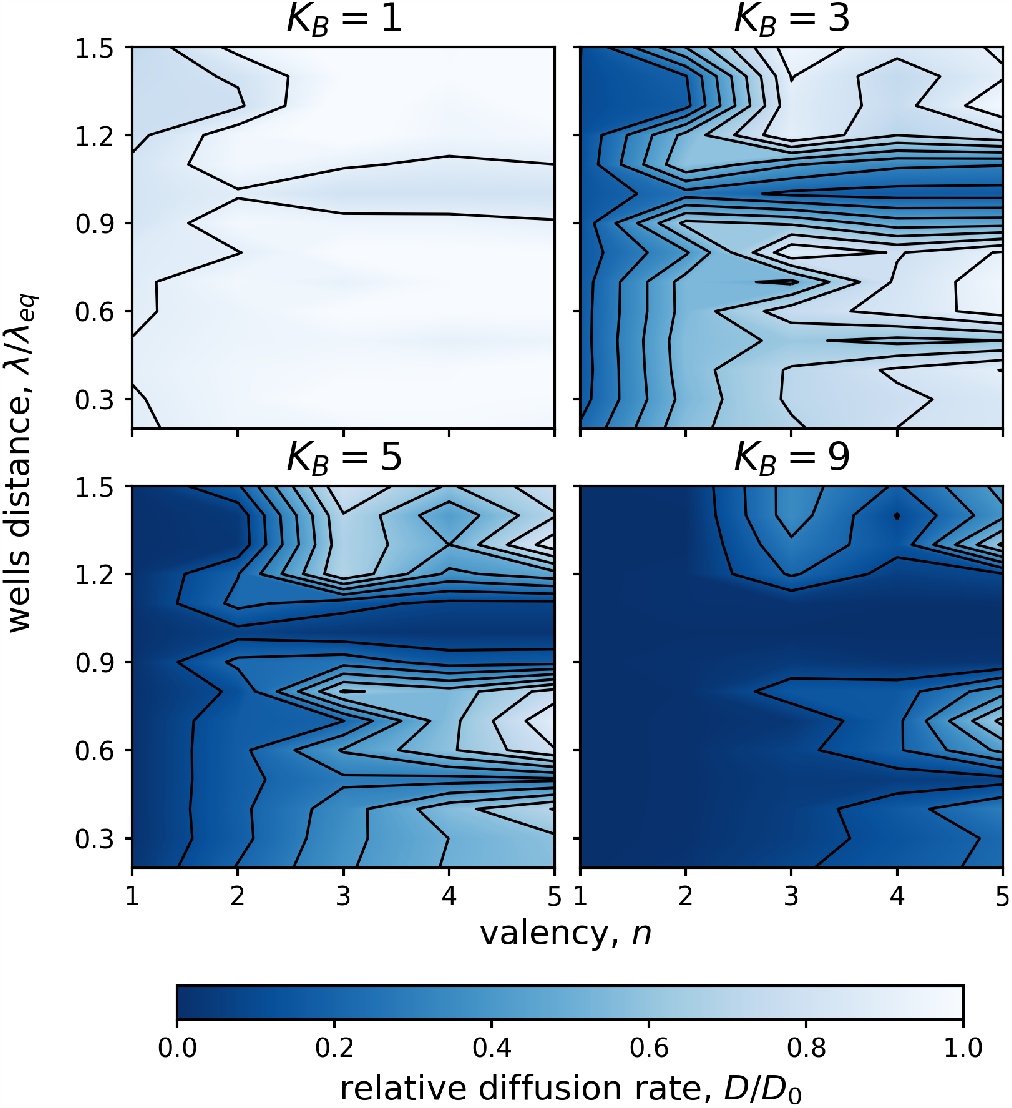
Dependence of the 1D relative diffusion rate, *D/D*_0_, on valency, *n*, and wells distance, *λ/λ*_eq_. Four net surface affinities, *K*_*B*_*∈ {*1, 3, 5, 9 *}*, were used. All ligands were connected with stiff linkers (*k* = 50). Note that if the wells distance matched the linker length *λ/λ*_eq_ *≈*1.0, the relative diffusion rate was independent of the valency. Similar, but weaker, effect was observed also for *λ/λ*_eq_ *≈* 0.5.

Finally, we show that the observed increase of the relative diffusion rate with increasing valency is independent of the distribution of binding and non-binding ligands within the particles (see Figure S4). However, the exact values of the relative diffusion rates are affected by the distribution of the binding ligands (see Figure S5).

## Discussion

We conducted dynamic Monte Carlo simulations with a simple model of multivalent particles to investigate their diffusion on a molecular surface and tether. Specifically, we focused on the scenario where multivalent particles bind with varying number of binding ligands to a host surface covered with fixed receptors sites and move laterally without rolling, i.e. walking is represented by binding and unbinding of the individual ligands to the receptors. The other mechanisms of diffusion, such as particle rolling, sliding along with attached receptors, and unbinding/rebinding with diffusion in solution have previously been described, ^28–32^ but their contribution to diffusion extends beyond the scope of this study.

Our results reveal that particles can have dramatically enhanced diffusion with increasing valency. This observation may seem counterintuitive because it has been established that more bonds between the particle and the host (i.e., increasing valency) leads to a reduction in the diffusion rate.^33–35^ However, in that case the strength of ligand-receptor bonds remained the same and increasing valency resulted in overall much stronger particle-host adsorption. In our model, we focused on the particles with the same binding affinity to the host (tether or surface). In particular, we decreased the strength of individual ligand-receptor bonds with increasing valency (keeping the valency times ligand-receptor interaction strength constant), which facilitated unbinding of the individual ligands and enabled particle walking. Our findings are thus in line with the reported faster diffusion rate of particles with weaker individual bonds between the ligand and receptors.^36,37^ Moreover, prior research has shown that continuous motion of motor proteins along microtubules relies on multivalency and the individual binding and unbinding of domains.^38^

In general, the net particle-host affinity could differ from the calculated multiplication of valency and the ligand-receptor binding strength because of possible combinations between bound receptors and ligands.^15^ Our particle-host affinity is thus a lower estimate for the given valency and multivalent particles with the same diffusion rate we obtained could bind to the host stronger in reality. Therefore, multivalent particles could diffuse faster despite having stronger affinity to the host.

Additionally, our model predicts that the diffusion rate is significantly reduced for particles with specific pattern/distribution of ligands. This reduction occurs when the ligand pattern matches the pattern of the receptors and the patterns are stiff. In such cases, all binding ligands align with the receptors, resulting in diminished diffusion. This “pattern matching” effect is more pronounced for stiffer patterns and higher valencies, while flexible patterns remain unaffected. Based on these results we anticipate scenarios where the host bound particles exhibit rapid diffusion on surface with heterogeneously distributed receptors until these particles find a matching receptor pattern where the particles are likely to halt.

In summary, we demonstrated that multivalent particles can achieve remarkable diffusion acceleration with increasing valency, while bound to a host tether or a surface. By distributing the binding affinity to multiple ligands, the initially non-diffusing monovalent particle could achieve a diffusion rate nearly as high as that of an unbound particle. The fast diffusion could be controlled for particles which have rigid distribution of ligands that matches the distribution of host receptors. Our results could find applications in various fields utilizing the design of fast-diffusing particles that maintain a strong affinity for target, such as molecular walkers^39^ or DNA-binding sliding proteins.^40–42^

## Acknowledgement

This work was supported by the European Research Council (ERC) under the European Union’s Horizon 2020 research and innovation programme (grant agreement No 101001470) and the project National Institute of virology and bacteriology (Programme EXCELES, ID Project No. LX22NPO5103) - Funded by the European Union - Next Generation EU. For financial support, ML thanks the Swedish Research Foundation and the Crafoord Foundation. Computational resources were provided by the CESNET, CERIT Scientific Cloud, and IT4 Innovations National Supercomputing Center by MEYS CR through the e-INFRA CZ (ID:90254). We acknowledge curated use of large language models (ChatGPT) for linguistic modifications of the article.

## Supporting Information Available

Supporting Information is available free of charge at XXXXX. All simulation data are provided at doi.org/10.5281/zenodo.8396688. The code used to perform the simulations is available from doi.org/10.5281/zenodo.8340209.

## Supporting Information

### Simulated distributions of interacting ligands

**Table S1:**
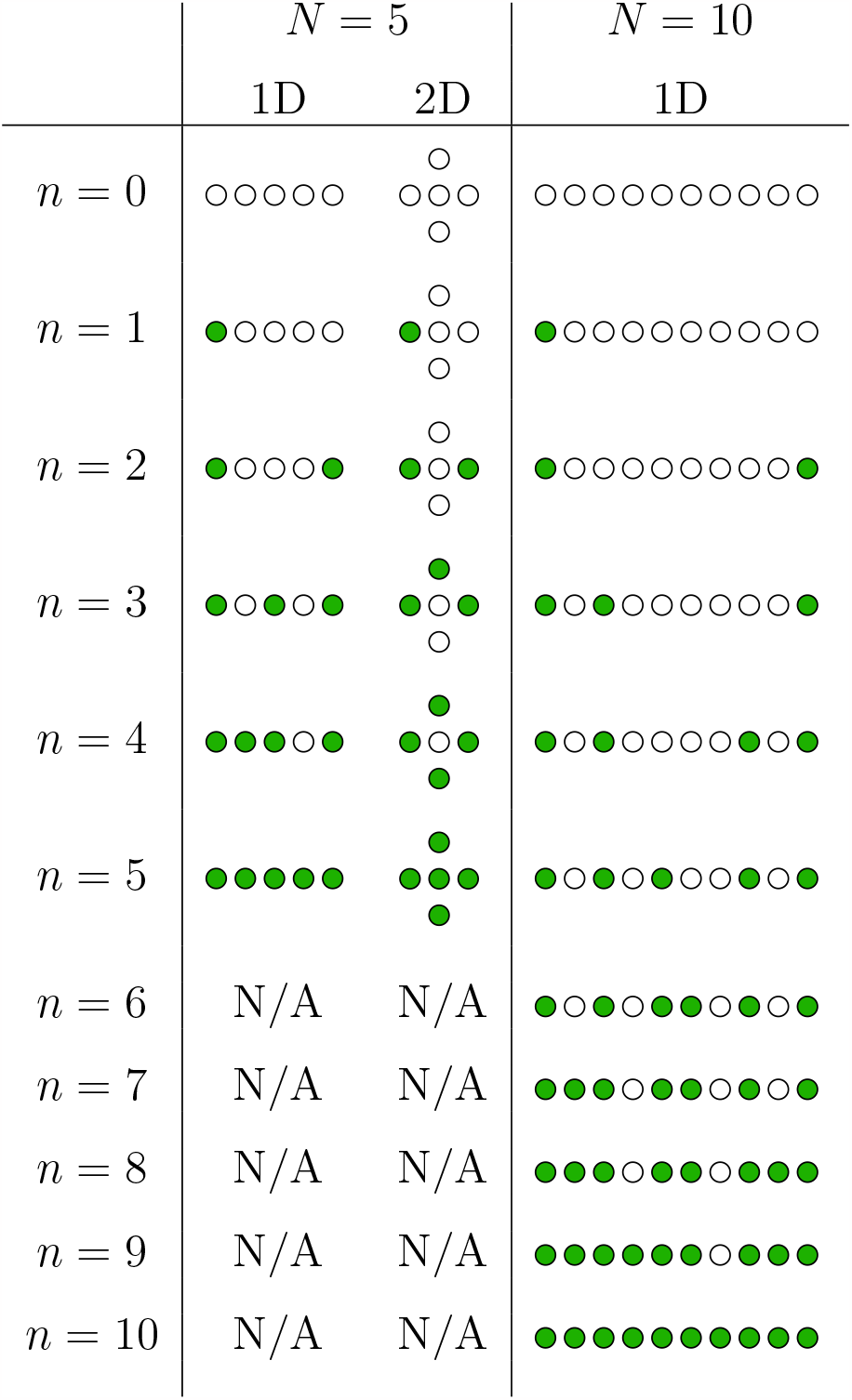
Standard, quasi-homogeneous distribution of ligands used for the simulations presented in the main text, unless stated otherwise. 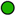 corresponds to a ligand that interacts with the surface potential (binds to receptors), while ◯ corresponds to a non-binding ligand.

**Table S2:** Distributions of ligands used for 2D simulations presented in Figure S4. 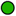 corresponds to a ligand that interacts with the surface potential (binds to receptors), while ◯ corresponds to a non-binding ligand.

|  | linear chain | alternative star-shaped |
| --- | --- | --- |
| $n = 0$ | ○○○○○ | ○<br>○○○<br>○ |
| $n = 1$ | ●○○○○ | ○<br>○●○<br>○ |
| $n = 2$ | ●○○○● | ○<br>●●○<br>○ |
| $n = 3$ | ●○●○● | ○<br>●●●<br>○ |
| $n = 4$ | ●●●○● | ●<br>●●●<br>○ |
| $n = 5$ | ●●●●● | ●<br>●●●<br>● |

Figure S5 (at the end of the Supporting Information) shows results obtained for the quasi-homogeneous pattern 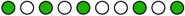 and for the concentrated pattern 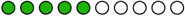.

### Example of mean-square deviation curves

**Figure S1:**
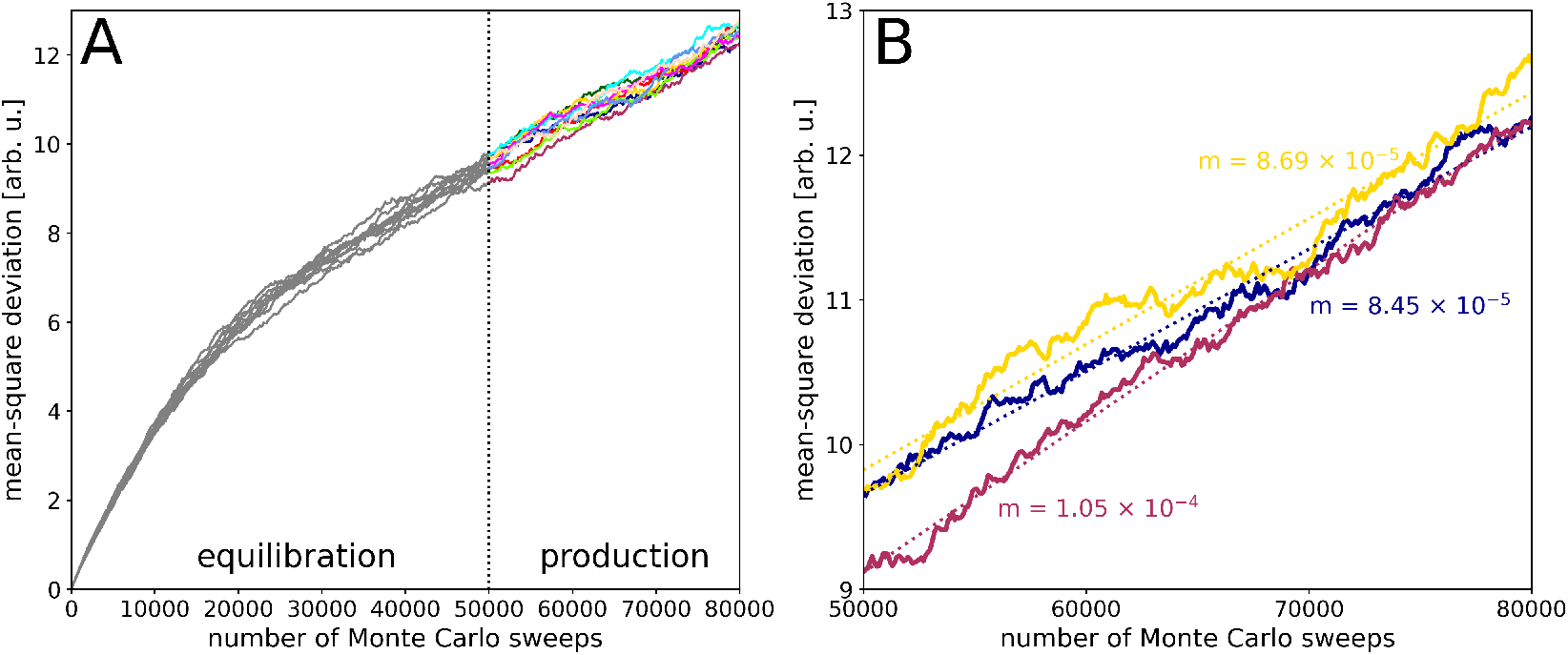
A. Ten mean-square deviation (MSD) curves collected over 30,000 independent simulations of a particle with *N* = 5, *n* = 1, *k* = 1, *λ/λ*_eq_ = 1.0, *K*_*B*_ = 3.0 diffusing on a 1D line. MSD data were collected over the entire simulation run, but data from the first 50,000 sweeps of each replica (equilibration stage) were not used for line fitting. B. Three selected MSD curves obtained from the same system, displaying fitted lines and their slopes during the production phase of the simulation. The diffusion rate for each curve was calculated by dividing the slope by a scaling factor based on the dimensionality of the problem (2 for 1D or 4 for 2D) and multiplying by an arbitrary constant (1000). The diffusion rates from individual MSD curves were averaged, and their standard deviation represents the error estimate. Throughout the main text, diffusion rates are reported relative to free diffusion calculated using the same method. The line fitting utilized the Rust [linreg](https://crates.io/crates/linreg) crate v0.2.0, while the mean and standard deviation calculations were conducted using the [statistical](https://crates.io/crates/statistical) crate v1.0.0.

### Diffusion rate

**Figure S2:**
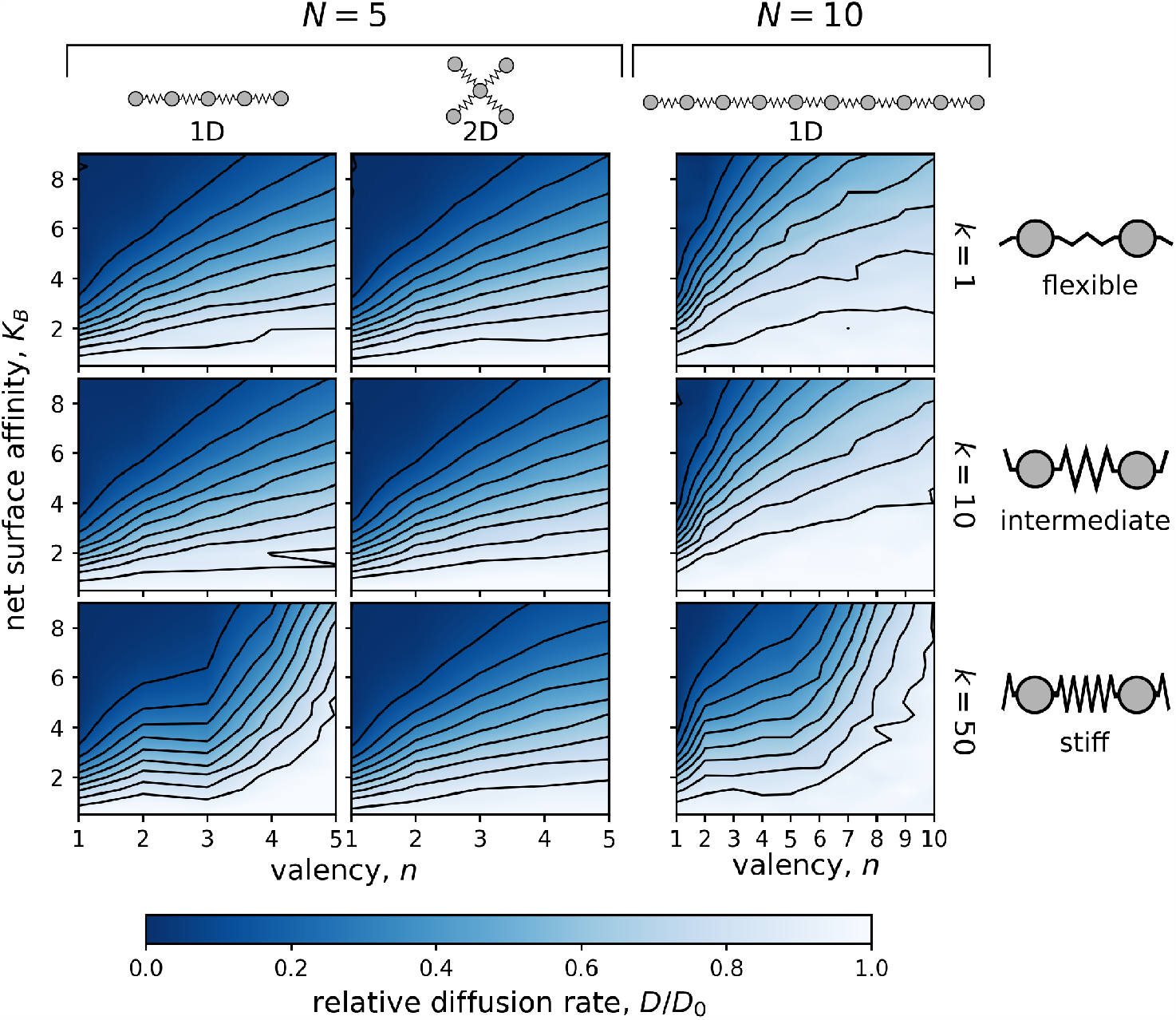
Dependence of the relative diffusion rate, *D/D*_0_ on the valency, *n*, and the net surface affinity, *K*_*B*_. Simulations were performed using particles consisting of either *N* = 5 (two columns on the left) or *N* = 10 ligands (column on the right) connected by flexible (*k* = 1; top row), intermediate (*k* = 10; middle row), or stiff (*k* = 50; bottom row) linkers. The distance between the surface potential wells, *λ/λ*_eq_, was set to 0.7. Diffusion of particles with *N* = 5 was calculated in both 1D (linear chain) and 2D (star) geometries, while diffusion of particles with *N* = 10 was examined only in the 1D case (linear chain). Note that for stiffly linked ligands (*k* = 50), the 1D relative diffusion rate increases exponentially with increasing valency. This behavior is in contrast to what was observed in Figure 3 and it is the consequence of the wells distances not matching the length of the linkers (*λ/λ*_eq_ = 0.7).

### Detailed analysis of pattern matching

As mentioned in the main text, we observed the lowest diffusion rate for particles whose linkers were slightly *shorter* than the distance between the surface potential wells. Specifically, the relative diffusion rate, represented by *D/D*_0_, was found to be lower for systems with *λ/λ*_eq_ = 1.1 compared to systems with *λ/λ*_eq_ = 1.0. This trend was consistently observed across a wider range of particles with *N ∈{* 2 … 5*}* and *k* = 10 diffusing on a line (1D), as depicted in Figure S3 A.

For further analysis, we focused on the simplest case of a two-binding ligand particle, which was diffusing on a line (1D). In this case, the unnormalized (absolute) mean diffusion rates *D* were calculated as 0.176 *±*0.011 and 0.157 *±*0.011 arb.u. for *λ/λ*_eq_ = 1.0 and *λ/λ*_eq_ = 1.1, respectively. To gain insights into the underlying mechanisms, we calculated the potential energy surfaces for these two systems, considering the two degrees of freedom of the two-ligand particle, as depicted in Figure S3 B. Both potential energy surfaces exhibited characteristic diagonally repeating local minima. Surprisingly, as shown in S3 C, the potential energy barriers between these minima were lower for the system with slower diffusion (*λ/λ*_eq_ = 1.1), suggesting that the observed differences in diffusion rates were originating from entropic effects. Our hypothesis was that the faster diffusion for *λ/λ*_eq_ = 1.0 was due to the presence of four distinct paths that the ligand could take to transition from one local energy minimum to a neighboring minimum, as indicated by the white arrows in Figure S3 B. In contrast, the potential energy surface for *λ/λ*_eq_ = 1.1 only presented two energetically favorable paths that allowed the ligand to leave each minimum.

To test this hypothesis, we conducted additional simulations for each of the described systems (with *λ/λ*_eq_ values of 1.0 and 1.1). In these simulations, we implemented a restriction to prevent the ligands from entering specific regions of the configuration space, as indicated by the white circles in S3 B. To achieve this, we applied a restraint in the following form:

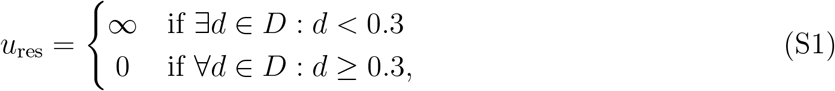

where *D* is a set of distances between the ligands’ coordinates *x*_*i*_ and *x*_*j*_ and the centers of the restricted areas, defined as

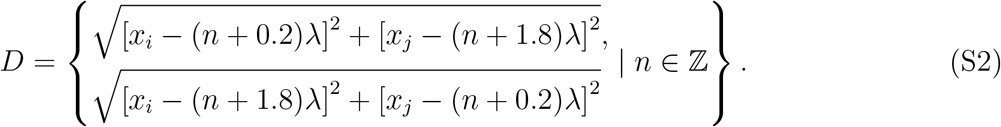

The applied restraint had a significant impact on the diffusion behavior of the particle with *λ/λ*_eq_ = 1.0. By obstructing half of the energetically most favorable paths, the diffusion rate of the particle was reduced by approximately 40% to 0.104 *±* 0.008 arb.u. In contrast, the effect of the restraint on the system with *λ/λ*_eq_ = 1.1 was milder. It led to a modest decrease in the diffusion rate of about 16% to 0.132 *±*0.007 arb. u., as the particle already exhibited a preference for the unblocked paths.

This showed that if the same number of paths was available for the particle to take, the diffusion rates were consistent with the height of the potential energy barriers (i.e. lower potential energy barriers between the minima led to higher diffusion rate) and that the originally observed discrepancy was indeed introduced by entropic effects.

In the next two paragraphs, we attempt to provide an intuitive explanation for the existence of 4 or 2 energetically favorable paths. When the distance between the wells perfectly matched the length of the linker (*λ/λ*_eq_ = 1.0) and both ligands of the particle resided in neighboring potential wells, the linker would be in equilibrium, meaning there would be no tension in the linker. In such case, the particle’s diffusion could be initiated by either ligand moving in either direction, resulting in the stretching or compressing of the linker. Thus, there were four possible ways for the particle to leave the energy minimum, as depicted in the potential energy surface for *λ/λ*_eq_ = 1.0.

**Figure S3:**
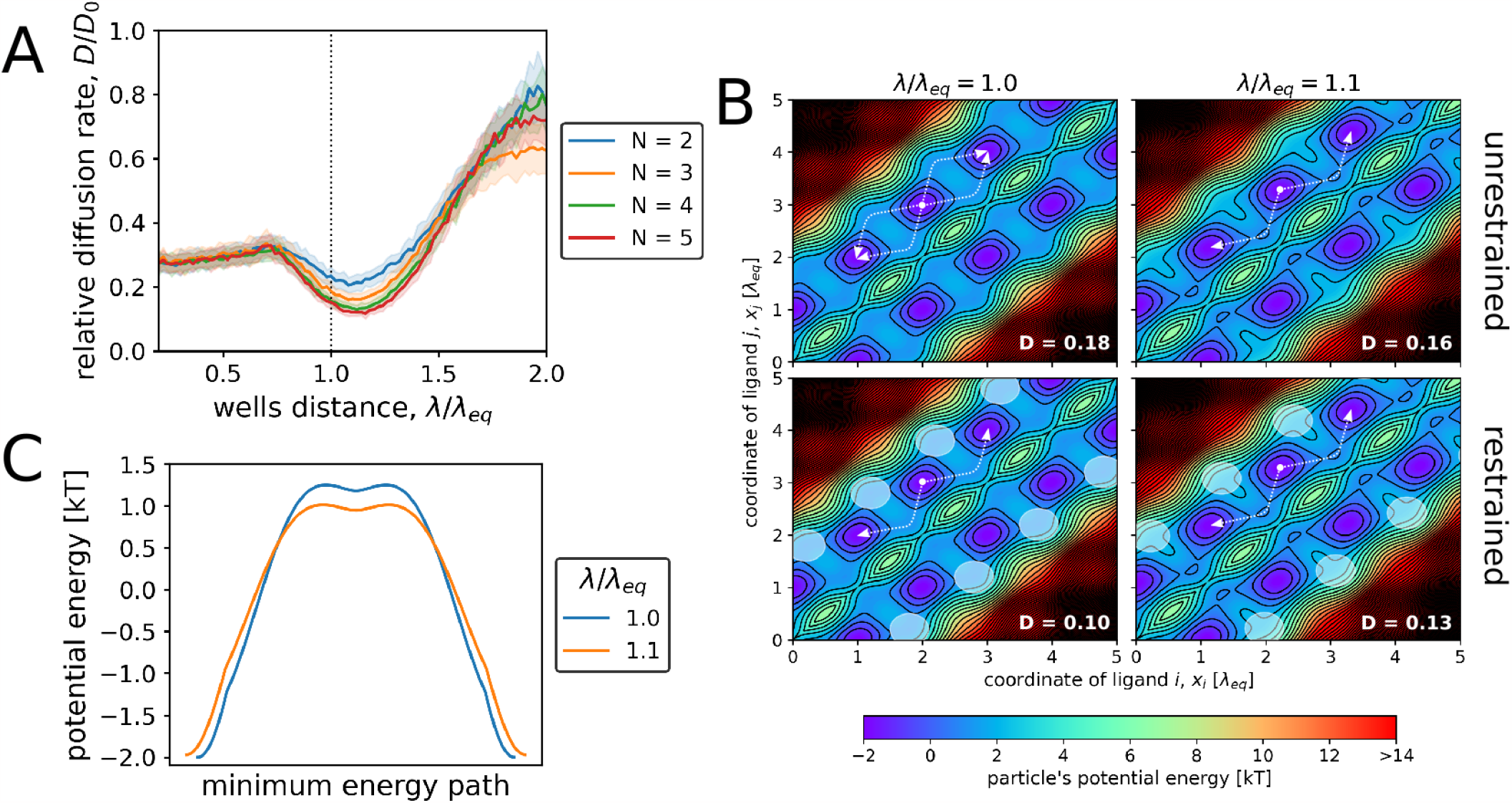
A. Dependence of 1D relative diffusion rate, *D/D*_0_, on the distance between the surface potential wells, *λ/λ*_eq_, for particles composed of varying number of ligands, *N*. All ligands were binding (*n* = *N*) and all ligands were connected into a linear chain using linkers with a force constant, *k*, of 10. B. Slices of potential energy surfaces for a particle composed of two intermediately linked (k = 10) binding ligands in a system with *λ/λ*_eq_ = 1.0 (left column) or *λ/λ*_eq_ = 1.1 (right column). Potential energy is shown as a function of the coordinates of the particle’s ligands. The white arrows show energetically favorable paths the particle can take to move between neighbouring potential energy minima. In the bottom row, the white circles show areas that the particle was restricted from in the restrained simulations. In the lower right corner of each chart, we show the unnormalized diffusion rate calculated for this potential energy surface (in arb. u.). C. Potential energy profiles of the particle along the identified paths between two neighbouring energy minima, calculated for *λ/λ*_eq_ = 1.0 and for *λ/λ*_eq_ = 1.1. The energies were obtained from the analytically calculated potential energy surfaces using the MEPSAnd script version 1.6 (available from bioweb.cbm.uam.es/software/MEPSAnd).

In contrast, if the linker was slightly shorter than the distance between the wells (*λ/λ*_eq_ = 1.1) and both ligands of the particle were located in neighboring potential wells, the linker was already slightly stretched, creating tension that forced the ligands closer together. Consequently, in such scenario, the particle’s diffusion could not be initiated by either ligand moving *away* from the other, as this would involve energetically unfavorable stretching of the already stretched linker. Instead, the process of the particle leaving the energy minimum had to be triggered by either of the ligands moving *closer* to the other, bringing the linker closer to equilibrium. Therefore, the number of available paths is reduced to two, as illustrated in the potential energy surface for *λ/λ*_eq_ = 1.1.

A similar mechanism is likely to apply also to particles composed of a larger number of ligands, which could explain the unexpected slowdown in diffusion at *λ/λ*_eq_ = 1.1.

### Effect of ligands distribution on diffusion

**Figure S4:**
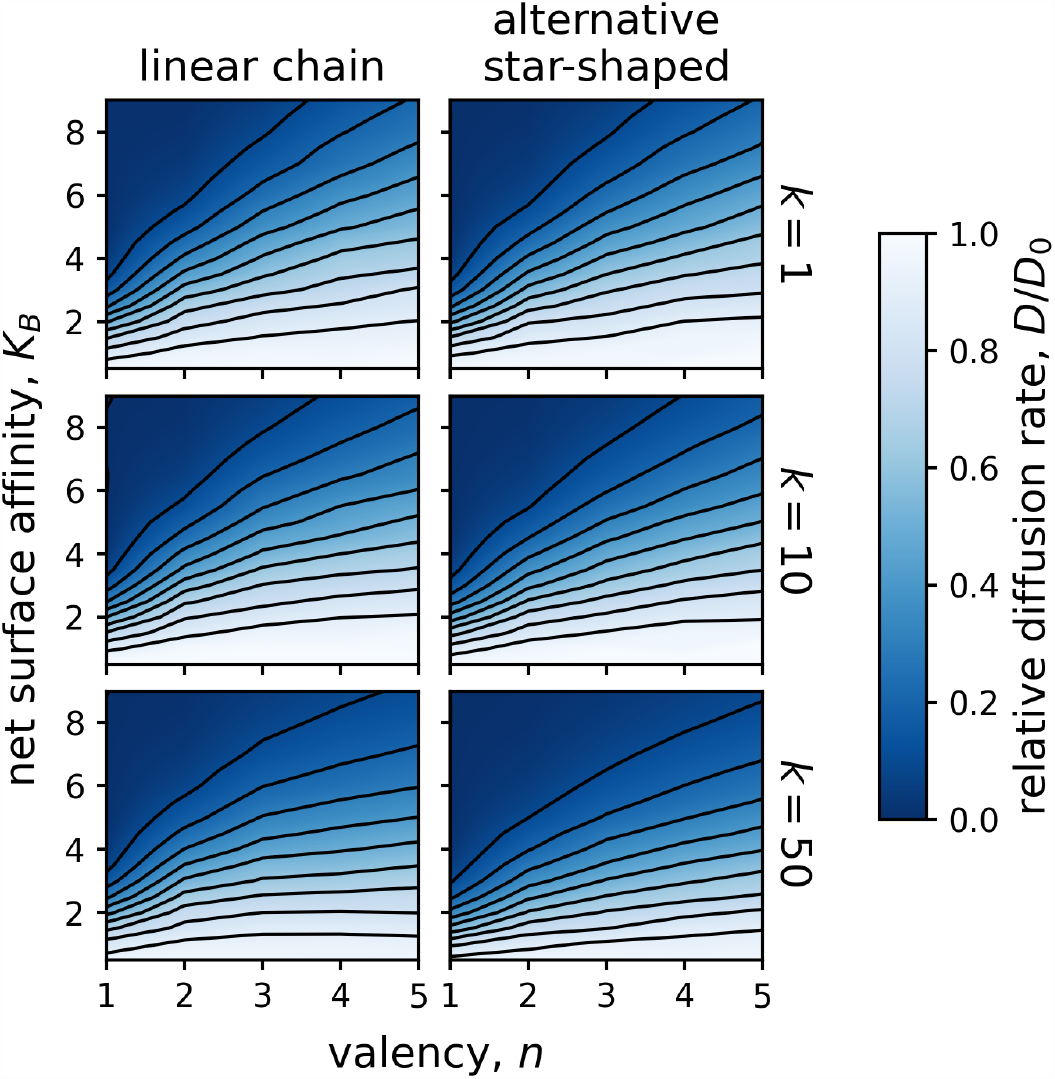
Dependence of the relative diffusion rate, *D/D*_0_, on the valency, *n*, and the net surface affinity, *K*_*B*_, for *N* = 5 particles with different geometries diffusing on a plane (2D). The left column shows results for particles with ligands arranged in a linear chain, while the right column shows results for alternative star-shaped particles, where the central ligand always interacted with the surface potential (unless *n* = 0). Note that for the standard star-shaped particles, the central ligand was always non-binding, unless *n* = *N*. See Table S2 for a detailed depiction of the geometries used in these simulations. The simulated particles consisted of flexibly (*k* = 1; top row), intermediately (*k* = 10; middle row), or stiffly linked ligands (*k* = 50; bottom row), and the distance between the surface potential wells matched the equilibrium linker lengths. Note that the overall trends in the diffusion rates matched the trends described in the main text for the standard distributions of the binding ligands.

**Figure S5:**
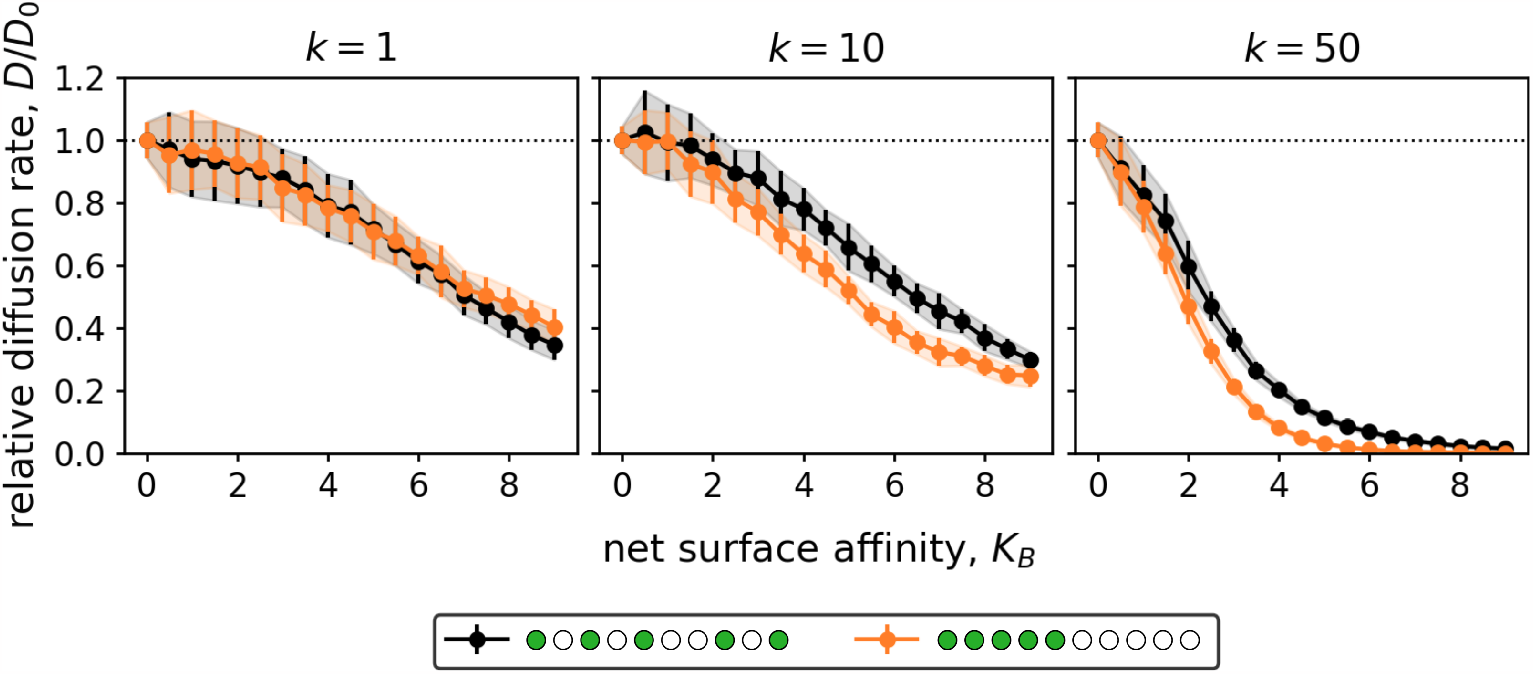
The effect of binding ligand distribution on 1D diffusion. Particles consisting of *N* = 10 ligands, 5 of which were binding (*n* = 5), connected by flexible (*k* = 1), intermediate (*k* = 10), or stiff (*k* = 50) linkers were studied. The distance between the surface potential wells matched the equilibrium linker lengths. The black lines represent the quasi-homogeneous distribution of binding ligands used for most simulations, while the orange lines correspond to the alternative distribution in which all binding ligands were concentrated at one end of the chain. Note that for sufficiently tightly linked particles (*k≥* 10), the concentrated pattern of binding ligands led to slightly lower diffusion rates than the quasi-homogeneous pattern. This was not observed for particles with *k* = 1 as these ligands were so weakly linked that they could exchange their positions in the chain.

## Notes

### Competing Interest Statement

The authors have declared no competing interest.

### Summary of Updates

References, author orcid

## References

(1) Lee, R. T., Ichikawa, Y., Kawasaki, T., Drickamer, K., Lee, Y. C. Multivalent lig- and binding by serum mannose-binding protein. Archives of Biochemistry and Biophysics 1992, 299, 129–136.

(2) Mondal, S., Narayan, K., Botterbusch, S., Powers, I., Zheng, J., James, H. P., Jin, R., Baumgart, T. Multivalent interactions between molecular components involved in fast endophilin mediated endocytosis drive protein phase separation. Nature Communications 2022, 13, 5017.

(3) Banjade, S., Rosen, M. K. Phase transitions of multivalent proteins can promote clustering of membrane receptors. eLife 2014, 3, e04123.

(4) Cuesta, A. M., Sainz-Pastor, N., Bonet, J., Oliva, B., Alvarez-Vallina, L. Multivalent antibodies: when design surpasses evolution. Trends in Biotechnology 2010, 28, 355–362.

(5) Dam, T. K., Roy, R., Das, S. K., Oscarson, S., Brewer, C. Binding of Multivalent Carbohydrates to Concanavalin A andDioclea grandiflora Lectin. Journal of Biological Chemistry 2000, 275, 14223–14230.

(6) Lu, W., Pieters, R. J. Carbohydrate–protein interactions and multivalency: implications for the inhibition of influenza A virus infections. Expert Opinion on Drug Discovery 2019, 14, 387–395.

(7) Müller, C., Despras, G., Lindhorst, T. K. Organizing multivalency in carbohydrate recognition. Chem. Soc. Rev. 2016, 45, 3275–3302.

(8) Overeem, N. J., Vries, E., Huskens, J. A Dynamic, Supramolecular View on the Multivalent Interaction between Influenza Virus and Host Cell. Small 2021, 17, 2007214.

(9) Delguste, M., Zeippen, C., Machiels, B., Mast, J., Gillet, L., Alsteens, D. Multivalent binding of herpesvirus to living cells is tightly regulated during infection. Science Advances 2018, 4, eaat1273.

(10) Paran, N. HBV infection of cell culture: evidence for multivalent and cooperative attachment. The EMBO Journal 2001, 20, 4443–4453.

(11) Martinez-Veracoechea, F. J., Mognetti, B. M., Angioletti-Uberti, S., Varilly, P., Frenkel, D., Dobnikar, J. Designing stimulus-sensitive colloidal walkers. Soft Matter 2014, 10, 3463–3470.

(12) Van Dongen, M. A., Dougherty, C. A., Banaszak Holl, M. M. Multivalent Polymers for Drug Delivery and Imaging: The Challenges of Conjugation. Biomacromolecules 2014, 15, 3215–3234.

(13) Woythe, L., Tito, N. B., Albertazzi, L. A quantitative view on multivalent nanomedicine targeting. Advanced Drug Delivery Reviews 2021, 169, 1–21.

(14) Zhang, Y., Yu, Y., Jiang, Z., Xu, H., Wang, Z., Zhang, X., Oda, M., Ishizuka, T., Jiang, D., Chi, L. et al. Single-Molecule Study on Intermolecular Interaction between C _60_ and Porphyrin Derivatives: Toward Understanding the Strength of the Multivalency. Langmuir 2009, 25, 6627–6632.

(15) Martinez-Veracoechea, F. J., Frenkel, D. Designing super selectivity in multivalent nano-particle binding. Proceedings of the National Academy of Sciences 2011, 108, 10963–10968.

(16) Leckband, D., Sivasankar, S. Mechanism of homophilic cadherin adhesion. Current Opinion in Cell Biology 2000, 12, 587–592.

(17) Sauer, M. M., Jakob, R. P., Luber, T., Canonica, F., Navarra, G., Ernst, B., Unverzagt, C., Maier, T., Glockshuber, R. Binding of the Bacterial Adhesin FimH to Its Natural, Multivalent High-Mannose Type Glycan Targets. Journal of the American Chemical Society 2019, 141, 936–944.

(18) Groves, J. T. Molecular Organization and Signal Transduction at Intermembrane Junctions. Angewandte Chemie International Edition 2005, 44, 3524–3538.

(19) Heldin, C.-H. Dimerization of cell surface receptors in signal transduction. Cell 1995, 80, 213–223.

(20) Cochran, J. R., Aivazian, D., Cameron, T. O., Stern, L. J. Receptor clustering and transmembrane signaling in T cells. Trends in Biochemical Sciences 2001, 26, 304–310.

(21) Puffer, E. B., Pontrello, J. K., Hollenbeck, J. J., Kink, J. A., Kiessling, L. L. Activating B Cell Signaling with Defined Multivalent Ligands. ACS Chemical Biology 2007, 2, 252–262.

(22) Block, S. M. Kinesin Motor Mechanics: Binding, Stepping, Tracking, Gating, and Limping. Biophysical Journal 2007, 92, 2986–2995.

(23) Hammer, J. A., Sellers, J. R. Walking to work: roles for class V myosins as cargo transporters. Nature Reviews Molecular Cell Biology 2012, 13, 13–26.

(24) Von Delius, M., Leigh, D. A. Walking molecules. Chemical Society Reviews 2011, 40, 3656.

(25) Valero, J., Škugor, M. Mechanisms, Methods of Tracking and Applications of DNA Walkers: A Review. ChemPhysChem 2020, 21, 1971–1988.

(26) Song, L., Zhuge, Y., Zuo, X., Li, M., Wang, F. DNA Walkers for Biosensing Development. Advanced Science 2022, 9, 2200327.

(27) Metropolis, N., Rosenbluth, A. W., Rosenbluth, M. N., Teller, A. H., Teller, E. Equation of State Calculations by Fast Computing Machines. The Journal of Chemical Physics 1953, 21, 1087–1092.

(28) Lee-Thorp, J. P., Holmes-Cerfon, M. Modeling the relative dynamics of DNA-coated colloids. Soft Matter 2018, 14, 8147–8159.

(29) Merminod, S., Edison, J. R., Fang, H., Hagan, M. F., Rogers, W. B. Avidity and surface mobility in multivalent lig- and–receptor binding. Nanoscale 2021, 13, 12602–12612.

(30) Evans, E., Sackmann, E. Translational and rotational drag coefficients for a disk moving in a liquid membrane associated with a rigid substrate. Journal of Fluid Mechanics 1988, 194, 553.

(31) Perl, A., Gomez-Casado, A., Thompson, D., Dam, H. H., Jonkheijm, P., Reinhoudt, D. N., Huskens, J. Gradient-driven motion of multivalent ligand molecules along a surface functionalized with multiple receptors. Nature Chemistry 2011, 3, 317–322.

(32) Blainey, P. C., Van Oijen, A. M., Banerjee, A., Verdine, G. L., Xie, X. S. A baseexcision DNA-repair protein finds intrahelical lesion bases by fast sliding in contact with DNA. Proceedings of the National Academy of Sciences 2006, 103, 5752–5757.

(33) Block, S., Zhdanov, V. P., Höök, F. Quantification of Multivalent Interactions by Tracking Single Biological Nanoparticle Mobility on a Lipid Membrane. Nano Letters 2016, 16, 4382–4390.

(34) Knight, J. D., Lerner, M. G., Marcano-Velázquez, J. G., Pastor, R. W., Falke, J. Single Molecule Diffusion of Membrane-Bound Proteins: Window into Lipid Contacts and Bilayer Dynamics. Biophysical Journal 2010, 99, 2879–2887.

(35) Marbach, S., Zheng, J. A., Holmes-Cerfon, M. The nanocaterpillar’s random walk: diffusion with ligand–receptor contacts. Soft Matter 2022, 18, 3130–3146.

(36) Jana, P. K., Mognetti, B. M. Translational and rotational dynamics of colloidal particles interacting through reacting linkers. Physical Review E 2019, 100, 060601.

(37) Lowensohn, J., Stevens, L., Goldstein, D., Mognetti, B. M. Sliding across a surface: Particles with fixed and mobile ligands. The Journal of Chemical Physics 2022, 156, 164902.

(38) Braun, M., Diez, S., Lansky, Z. Cytoskeletal organization through multivalent interactions. Journal of Cell Science 2020, 133, jcs234393.

(39) Leigh, D. A., Lewandowska, U., Lewandowski, B., Wilson, M. R. Synthetic Molecular Walkers. 2014, 354, 111–138, Series Title: Topics in Current Chemistry.

(40) Park, S., Lee, O.-c., Durang, X., Jeon, J.-H. A mini-review of the diffusion dynamics of DNA-binding proteins: experiments and models. Journal of the Korean Physical Society 2021, 78, 408–426.

(41) Givaty, O., Levy, Y. Protein Sliding along DNA: Dynamics and Structural Characterization. Journal of Molecular Biology 2009, 385, 1087–1097.

(42) Halford, S. E. How do site-specific DNA-binding proteins find their targets? Nucleic Acids Research 2004, 32, 3040–3052.

